# The effect of ocean warming on black sea bass (*Centropristis striata*) aerobic scope and hypoxia tolerance

**DOI:** 10.1101/507368

**Authors:** Emily Slesinger, Alyssa Andres, Rachael Young, Brad Seibel, Vincent Saba, Beth Phelan, John Rosendale, Daniel Wieczorek, Grace Saba

## Abstract

Over the last decade, ocean temperature in the U.S. Northeast Continental Shelf (U.S. NES) has warmed faster than the global average and is associated with observed distribution changes of the northern stock of black sea bass (*Centropristis striata*). Mechanistic models based on physiological responses to environmental conditions can improve future habitat suitability projections. We measured maximum, resting metabolic rate, and hypoxia tolerance (S_crit_) of the northern adult black sea bass stock to assess performance across the known temperature range of the species. A subset of individuals was held at 30°C for one month (30_chronic_°C) prior to experiments to test acclimation potential. Absolute aerobic scope (maximum – resting metabolic rate) reached a maximum of 367.21 mgO_2_ kg^−1^ hr^−1^ at 24.4°C while S_crit_ continued to increase in proportion to resting metabolic rate up to 30°C. The 30_chronic_°C group had a significant decrease in maximum metabolic rate and absolute aerobic scope but resting metabolic rate or S_crit_ were not affected. This suggests a decline in performance of oxygen demand processes (e.g. muscle contraction) beyond 24°C despite maintenance of oxygen supply. The Metabolic Index, calculated from S_crit_ as an estimate of potential aerobic scope, closely matched the measured factorial aerobic scope (maximum / resting metabolic rate) and declined with increasing temperature to a minimum below 3. This may represent a critical value for the species. Temperature in the U.S. NES is projected to increase above 24°C in the southern portion of the northern stock’s range. Therefore, these black sea bass will likely continue to shift north as the ocean continues to warm.

## Introduction

Marine environments are progressively warming as a consequence of climate change [1]. Along the U.S. Northeast Shelf (U.S. NES), ocean temperature is rising faster than the global average [2,3] resulting in a significant temperature increase [4,5]. Sea surface and bottom temperatures in the U.S. NES are projected to rise an additional 4.1°C and 5.0°C, respectively, along the U.S. NES [6,7]. Contemporary ocean warming in the U.S. NES has been associated with distribution shifts of many economically and ecologically important fish species both in latitude and/or depth [7–11], associated with tracking local climate velocities [12]. Understanding and projecting shifts in fish distribution will be important for characterizing potential ecological and economic impacts and anticipating and resolving fishery management conflicts [13].

Temperature directly affects metabolic rates of marine ectotherms [14,15] and is believed to set the boundaries of species ranges [16,17]. One explanation for the effects of temperature on ectothermic species, oxygen and capacity-limited thermal tolerance (OCLTT; [18]), postulates that thermal limitation occurs due to a mismatch in oxygen demand and supply at sub-optimal temperatures, which ultimately determines suitable thermal habitat [19]. In this framework, the thermal optimum occurs where absolute aerobic scope (AAS), the difference between maximum (MMR) and resting metabolic rate (RMR) [20], is highest. RMR is the cost of maintenance for an organism and increases with temperature [15]. MMR may be constrained differently across a temperature range by oxygen uptake, transport or utilization. The drop in AAS beyond the thermal optimum is associated with the failure of MMR to increase relative to the continuing rate of increase in RMR [21]. Absolute AAS is thought to represent the capacity for oxygen uptake, beyond that supporting maintenance metabolism, that can be utilized for activities that promote individual fitness (e.g. growth, reproduction, predator avoidance; [22]). Hence, the adaptive benefit of living at suitable temperatures to maintain metabolic scope may provide a mechanistic explanation for where fish may be distributed in their environment.

While the general distribution of fishes is broadly confined by thermal preferences, oxygen availability can further constrain suitable habitat. The hypoxia tolerance of a fish can be estimated as the critical oxygen saturation level (S_crit_), the %O_2_ below which oxygen supply cannot match the demands of maintenance metabolism. Further reductions in %O_2_ cause a proportional decrease in RMR [23]. Below the S_crit_, a fish has time-limited survival as ATP production progressively relies on unsustainable anaerobic pathways and metabolic suppression [24,25]. Generally, a fish with a low S_crit_ is tolerant of lower sustained oxygen levels [26]. As ocean temperature increases, oxygen demand concomitantly increases [27,28], potentially reducing hypoxia tolerance [29,30]. The S_crit_ further provides a means of calibrating the Metabolic Index, which is a ratio of oxygen supply to demand that provides an estimate of sustained factorial aerobic scope and metabolically suitable habitat [17]. At S_crit_, by definition, supply exactly matches demand allowing statistical estimation of the equations’ physiological parameters.

The northern stock of black sea bass (*Centropristis striata*) on the U.S. NES extends from Cape Hatteras to the Gulf of Maine and is centered in the Mid-Atlantic Bight (MAB; [31]). These fish seasonally migrate from the continental shelf edge in cooler months to inshore depths (5-50m) in warmer months [32,33]. Seasonally migrating black sea bass thus experience a wide range of temperatures throughout the year, ranging from 6°C during winter and up to 27°C during summer/early fall months [34]. Off the coast of New Jersey, periodic hypoxic events can occur during the summer as a result of high biological activity [35] fueled by upwelling of nutrient rich waters [36]. Therefore, during the warm summer months, black sea bass are potentially subject to hypoxia in this region, contributing to the oxygen limitation that contains suitable habitat.

The northern stock of black sea bass may already be exhibiting poleward shifts, likely due to ocean warming [7,37]. Evidence for current black sea bass distribution shifts comes primarily from bottom trawl survey data [7], and is supported anecdotally through fishermen. Laboratory-based process studies focused on the physiology of an organism provide detailed mechanistic relationships between the environment and the animal [38]. Results from these physiological studies are useful for modeling suitable habitat based on environmental parameters [39] and could be used to model black sea bass distributions (e.g., [17,40]), and project future distribution shifts in black sea bass with continued ocean warming. The objectives of this study were to determine the AAS and S_crit_ for the northern stock of adult black sea bass, parameters that can be used in future habitat suitability modeling. AAS and S_crit_ were measured over a range of temperatures similar to those experienced by black sea bass during their summer inshore residency to compare thermal optima, if present, and Metabolic Index against the known temperature range of the species. We show that, in support of recent northward population shifts, black sea bass are physiologically limited at higher temperatures possibly due to limitation of oxygen supply relative to demand for growth and reproduction. As such, black sea bass will likely continue to expand northward as the U.S. NES continues to warm.

## Materials and Methods

### Fish Collection and Husbandry

Adult black sea bass (*Centropristis striata*) from the northern stock (length = 221-398mm; weight = 193.7-700.4g) were collected off the coast of New Jersey, USA at depths of 15-20m in early June from Sea Girt and Manasquan Reefs by fish traps (2016), and from local reefs off Sandy Hook by hook-and-line (2017). Fish were housed in the NOAA James J. Howard Marine Laboratory, held at ambient temperature (22 ± 1°C) and salinity (26ppt), maintained at a natural photoperiod for New Jersey summer, and fed daily to satiation on a diet of sand lance and silversides for up to two months during the experimental trials. Water temperature and salinity was monitored daily using a YSI (Pro-30; Yellow Springs, Ohio, USA), and water chemistry remained at suitable levels (< 20 uM nitrate, undetectable nitrite, < 0.05 uM ammonia, pH range of 7.98-8.04). Fish were acclimated to captive conditions two weeks prior to the trials, after which all experimental fish ate regularly and were in good condition. Any fish exhibiting apparent health issues or stress (i.e. lack of appetite, difficulties with buoyancy or orientation) were not used in experiments. Fish that could not recover apparent health issues and exhibited extreme distress were euthanized with an overdose of MS-222 (250 mgL^−1^) (see Ethics Statement below). After acclimation, fish were measured for length (TL mm), weight (g), and tagged with individually numbered T-bar Floy tags inserted underneath the dorsal rays. For each temperature treatment, fish were acclimated at a rate of 2°C day^−1^ to reach experimental temperature, then held at the target treatment temperature for at least 48hr prior to the start of experiments. Fish were starved 48hr prior to the start of each experiment to eliminate effects of specific dynamic action (41). At the end of each experiment, all fish were euthanized with an overdose of MS-222 (250 mgL^−1^) and dissected to obtain final sex of each fish (see Ethics Statement below).

### Experimental Set-Up

Experimental tanks (1,200L) were filled with treated seawater from Sandy Hook Bay that continuously circulated through a closed system. Circulating seawater was treated using filters (sand and biological) and UV-light, and salinity was adjusted to mimic average summertime inshore NJ bottom water (32±1). Experimental temperatures were achieved using in-line chillers (Aqua Logic Delta Star; San Diego, California, USA) and/or titanium exchanger heaters (Innovative Heat Concepts, Homestead, Florida, USA), and maintained at ±1°C from target temperature.

Metabolic rates were measured using intermittent respirometry under the protocols outlined in Clark et al. [42] and Svendsen et al. [43]. Flow-through respirometers (13.5 liter volume; 23[H]x26[W]x37[L] cm plexiglass) were placed into the two experimental tanks (two respirometers per tank; four respirometers per trial). Flush pumps (Eheim Universal 600 l//h; Deizisau, Germany) connected to the respirometer were used to pull water from the surrounding temperature bath to replenish dissolved oxygen and eliminate metabolic waste buildup within the respirometer. The duration and timing of flushes set the intermittent cycles, which were controlled through a pre-determined time sequence using a DAQ-M instrument (Loligo Systems; Viborg, Denmark), and were set based on the trial temperature so that oxygen saturation was never below 75% [44]. For each closed measure period (when flush pumps were off), the rate of decline in dissolved oxygen concentration within the sealed chamber was used to calculate a mass specific rate of oxygen consumption, or metabolic rate (MO_2_: mgO_2_ kg^−1^ hr^−1^). A closed recirculation loop connected with a smaller pump (Eheim Universal 300 l/h; Deizisau, Germany) was also utilized to uniformly disperse dissolved oxygen within the chamber and provide waterflow across the oxygen dipping probe optical mini sensor (PreSens Pst3; Regensburg, Germany). Oxygen probes were calibrated in accordance with the supplier’s manual (Oxygen dipping probe PSt3, PreSens GmbH, Regensburg, Germany) and checked with a YSI (ProSolo ODO; Yellow Springs, Ohio, USA) that was calibrated in 100% and 0% oxygen saturation sample waters. Autoresp computer software (Loligo Systems; Viborg, Denmark) and a Witrox-4 instrument (Loligo Systems; Viborg, Denmark) were used to continuously monitor dissolved oxygen and temperature within the chamber over the course of the experiment.

For hypoxia experiments, intermittent respirometry was also used to avoid a CO_2_ and metabolite build up [45]. Each respirometer flush pump was connected to an external water bath that was filled with the same system water. Within the external water bath, a pump (Eheim Universal 1200 l/h; Deizisau, Germany) connected to a piece of Tygon tubing held an oxygen optode to monitor source O_2_ and served as a mixing device. Also within the external water bath, four small microdiffusers were connected to a N_2_ gas canister [23] to allow for diffusion of nitrogen gas into the external bath and subsequent displacement of O_2_ within the external water bath and control of environmental %O_2_ within the chambers over the course of the hypoxia experiment.

Background respiration was measured by taking background MO_2_ (MO_2br_) pre- and post-trial in empty chambers for ~1.5hr. A linear regression between pre- and post-MO_2br_ was used to apply a correction factor to each MO_2_ value recorded throughout an experiment.

Experiments were conducted at a range of temperatures (12, 17, 22, 24, 27 and 30°C). An additional subset of black sea bass were held at 30°C for one month to test acclimation potential (S1 Dataset). We used two different methods in an attempt to elicit maximum metabolic rate (MMR): exhaustive-chase and swim-flume. For the chase method, individual black sea bass (S1 Dataset) were placed in a 4ft-diameter chase tank filled with water from the experimental tanks. Fish were chased via tactile stimulation on the caudal tail and were determined exhausted when unresponsive to further tactile stimulation and air exposure. Fish were then immediately transferred to individual respirometers that were sealed within ~1 min from the end of the chase and remained in the metabolic chambers for ~23hr allowing for resting metabolic rate (RMR) measurement [46]. Once the experiment was finished, fish were either exercised in a swim-flume (Loligo Systems 90L; Viborg, Denmark) after a 24hr rest period or remained in the chamber for the hypoxia experiments. For the swim-flume, fish were tested using a sprint protocol.

Swimming speed was increased over a 5min period up to 0.95 BL s^−1^ with a flush pump on as the fish adjusted to the flume. After an adjustment time (~10 minutes), the speed was increased over a period of 5 minutes until the fish was sprinting (designated as >10 tail bursts during 30s intervals and an inability to maintain position in the working section without burst swimming). Once a fish was sprinting, the flush pump was turned off and the flume was sealed. Fish were held at their sprint speed for at least 10 minutes or until failure, determined when the fish rested at the backgrate for >10s. Aerobic scope was calculated in absolute (AAS = MMR-RMR) and factorial terms (FAS = MMR/RMR). An additional subset of black sea bass was held at 30°C for one month to test acclimation potential. In 2016, fish were only tested at 24, 27, and 30°C due to restrictions in maintaining temperatures. See S1 Full Dataset (and accompanying S1 File with metadata description) for sample size at each temperature.

### Critical %O_2_ determinations

Hypoxia (S_crit_) experiments were conducted on the last 4 fish of each temperature treatment trial. This allowed us to reliably use fish that were already acclimated to the respirometers and had reached RMR overnight. Starting with 100% dissolved oxygen (DO) saturation within the chambers, environmental %O_2_ was incrementally decreased by 10%. Three intermittent (flush, wait, measure) cycles were measured per DO level until S_crit_ was determined to have been reached, indicated by a substantial decline in fish metabolic rate or loss of equilibrium, and the experiment ended.

### Ethics statement

All animal work has been conducted according to relevant national and international guidelines. Fish were collected under New Jersey permits #1610 & #1717. Protocols for the treatment and euthanasia procedure of all animals reported here was approved by the Rutgers University Institutional Animal Care and Use Committee (IACUC) (Protocol number 15-054). Most personnel working with animals on this project had previous training in vertebrate physiology research and were familiar with the protocols. Those without previous experience trained by co-PIs Grace Saba and Brad Seibel and NOAA Fisheries personnel prior to and during the experiments. All efforts were made to ensure minimal pain and suffering. No endangered or protected species were involved.

Throughout the duration of the study, fish in holding tanks and in active experiments were monitored frequently throughout the day. Because of the different demands placed on fish in holding vs. an experiment, there were different monitoring protocols for each. For fish in holding tanks prior to, during, and between experimental trials, the amount each fish ate during feeding was recorded, experimental pre- and post-weights were taken to determine if weight loss occurred, and daily visual inspections conducted served as a monitoring system for the fish well-being. The fish were housed in groups, but there were separate holding tanks dedicated for any individuals that exhibited distress or abnormal behavior (e.g., rapid drop in metabolic rate, no physical response to human disturbance, abnormal swimming behavior, inability to maintain proper position or buoyancy). This allowed for more direct monitoring of behavior and feeding, and minimized the chance of infection to spread if the fish was in fact ill. All researchers were instructed to minimize their presence in the room to decrease disturbances to the fish. Tanks were cleaned daily to help maintain good water quality.

Monitoring was also conducted in all aspects of the experimental trials. First, after chasing (for MMR), fish were placed in individual flow-through respirometers. The water bath tanks that held the respirometers during experimental trials were covered to minimize distraction from the researchers. The respirometers were connected to an AutoResp system that monitored real-time metabolic measurements without physically or visually disturbing the fish. If abnormal patterns in metabolic measurements were observed (e.g., rapid drop in metabolic rate), the fish was visually inspected for signs of distress and, if observed, promptly removed and placed in a holding tank for additional monitoring. Second, some fish swam in a swim-flume after a minimum 48hr recovery period after the respirometry chamber. Fish that refused to swim or exhibited distress or abnormal behavior were removed from the flume, placed in a holding tank for additional monitoring, and the experiment ended. Third, during the hypoxia tolerance experiments, the oxygen content of the water was slowly decreased. Fish were constantly monitored throughout the entire experiment and if they showed any sign of distress or abnormal behavior, the experiment was ended for that fish, and the fish was transferred to a holding tank. For all fish, they were never held beneath their critical oxygen saturation level for more than 30 minutes. Once fish were placed back into fully oxygenated water, they immediately began swimming and exhibited normal behavior. Finally, a subset of fish were held at 30°C chronically for one month. While this is not a lethal temperature for black sea bass in acute settings, it is unknown their response to a chronic exposure. This trial treatment was necessary in the context of this study because there will be portions of their habitat that will reach this temperature in 80 years under our current emissions scenarios. The purpose of this trial was not to investigate the lethal temperature for black sea bass, but to test their acclimation potential at 30°C by comparing their metabolic rates with those in the shorter 30°C treatment. Because these fish were being held at elevated temperatures, they were monitored very frequently and any sign of distress or abnormal behavior in the holding tanks during acclimation, or during or between an experiment resulted in additional monitoring and prompt euthanasia if required.

Throughout the experiment, there were two instances when a fish would be euthanized. First, all experimental animals were euthanized at the end of the experiment for two reasons: 1) To obtain a final sex of the fish, which needs to be macroscopically determined through dissections in this sex-changing fish species, and 2) To prevent any potential spread of pathogens or infectious disease to natural populations that may have resulted from prolonged captivity in the laboratory (~2 months) and gone undetected. During the 2016 and 2017 summer study periods, 164 fish were used and, as explained above, all were euthanized at the end of the experiments.

Second, if a fish exhibited distress or abnormal behavior during the experimental trials, the experimental treatment was terminated and the fish was placed in a separate holding tank and monitored. If the fish was not able to recover from its symptoms within 3 hours, it was immediately euthanized. Recovery was determined by the fish maintaining a proper balanced position at the bottom of the tank and changing color to their tank surroundings. If the fish recovered to this point, then the final determination of recovery was determined by whether the fish would regularly eat food within one week. A total of eleven fish were euthanized before the end of experiments, and these mortalities were all associated with the high temperature (30°C) treatment, likely due to deterioration of metabolic and/or muscle function. These fish either did not respond normally to nets entering the water (i.e. a healthy fish will dart away from the net) or exhibited a sudden drop in metabolic rate during experimental trials. These fish were monitored at a higher frequently in a separate holding tank and when their condition did not improve within 3 hours, they were immediately euthanized. No fish died without euthanasia in this study.

All instances of euthanasia were performed by an overdose of MS-222 (tricaine methanosulfate), a common anesthetic and euthanasia drug for fish. Death was determined by an observed absence of opercular movement/gill pumping for at least 10 minutes. After the fish was determined as dead in the MS-222, it was then pithed to ensure death.

### Data Analysis

Fish MO_2_ was calculated via the AutoResp program from the slope of oxygen saturation decline during each closed measurement period. Validation of each MO_2_ value was conducted using R_2_ values from each measure period. MO_2_ measurements with R_2_ values < 0.9 were not used.

We report our baseline metabolic rate as resting (RMR) instead of standard (SMR) metabolic rate because the amount of time in the chamber was ~23 hours, which does not allow for determination of full diel cycles [46]. RMR was calculated from a truncated dataset without the first two hours of elevated MO_2_ values following exercise and by using the 20th quantile of the RMR data in the *calcSMR* package in R [46]. Briefly, a frequency distribution of MO_2_ values from the truncated data set was created and the values at the 20^th^ quantile were taken to calculate RMR. MMR in the chase protocol was defined as the highest MO_2_ value recorded during the trial and MMR was calculated for the duration of the sprint interval in the swim-flume. AAS was taken as the difference between MMR and RMR. There was a significant effect of mass on MO_2_ (F_1,117_ = 4.651; *P* < 0.05; Fig. 1). Therefore, the effect of temperature on MO_2_ was analyzed using a one-way ANCOVA with weight as a covariate. A Tukey’s HSD *post hoc* was used to determine significant pair-wise comparisons between temperatures. MO_2_ was adjusted for weight using the estimated marginal means from the ANCOVA centered around a fish mean weight of 346.9g. Because the average weight of fish in the 24°C treatment (253.9g) was lower than the mean weight of all other experimental fish, the adjusted MO_2_ for 24°C was slightly overestimated and had larger standard error for AAS and MMR. Curves for aerobic scope were modeled using a 3^rd^ degree polynomial fit and were used to estimate a thermal optimum (temperature at the highest AS). All graphs and results report the adjusted MO_2_ (MO_2adj_).

**Figure 1.**
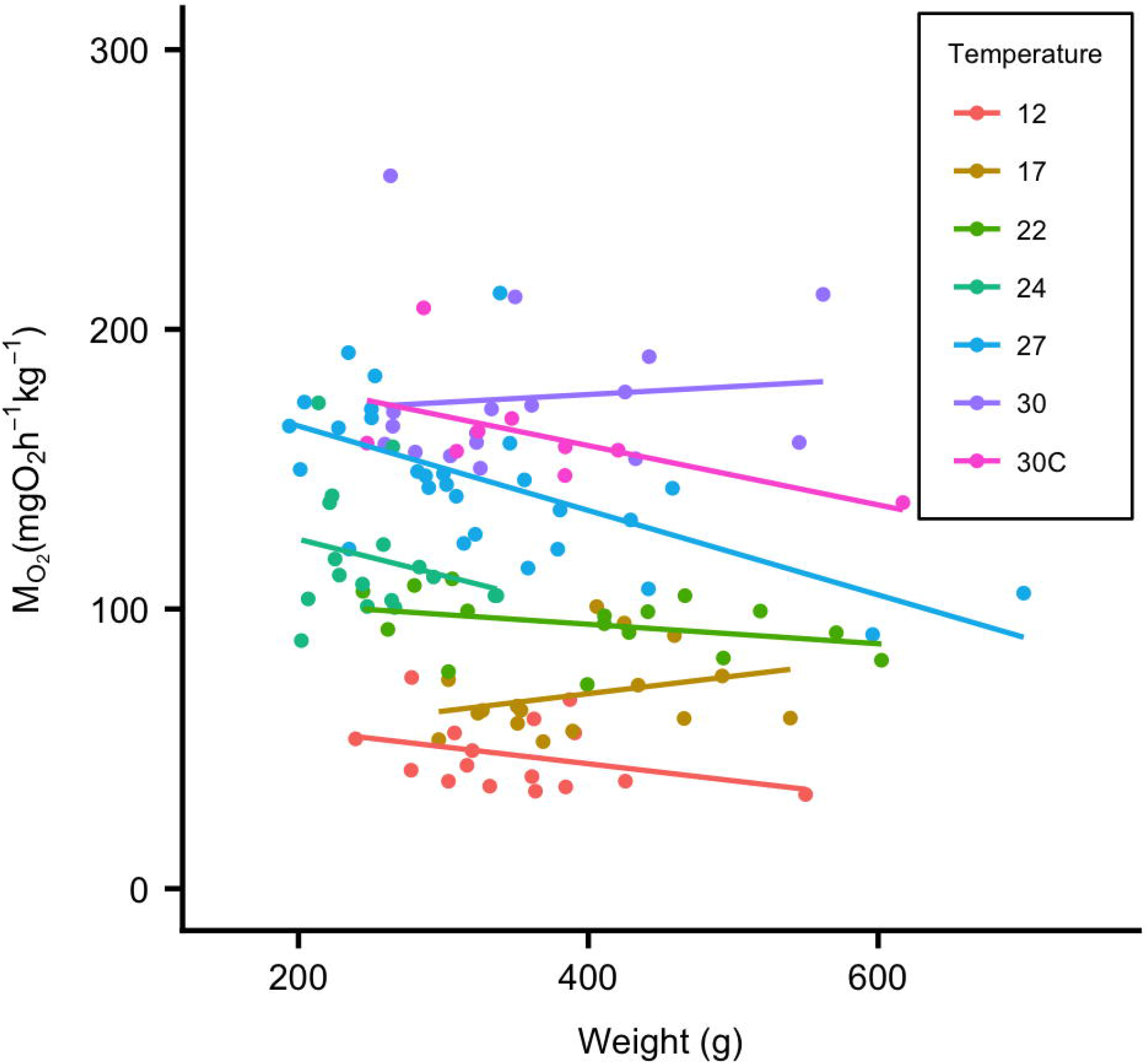
Temperature and body weight both affect resting metabolic rate in black sea bass. RMR (n=121) for each temperature treatment is plotted against body weight (g). A fitted regression line demonstrates that in addition to the effect of temperature on RMR, body weight also has an effect (*P*<0.05). 30c = 30_chronic_°C treatment.

Q_10_ values were calculated for the adjusted MO_2_ between temperature increments, and between the range of temperatures using the formula:

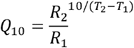

where Q_10_ is the temperature coefficient for MO_2_, R_1_ is the MO_2_ at T_1_ and R_2_ is the MO_2_ at T_2_. S_crit_ was determined by fitting two regression lines through the data: one through the region where RMR was independent of %O_2_ and one through the portion where MO_2_ decreased linearly with a decrease in %O_2_. The intersection of the two regression lines is the critical point used for S_crit_ [47]. This was analyzed using R code in the *calcO2crit* package from [26]. Because we had a sample size of 4 fish per temperature trial, a power analysis was run to determine the statistical power of this small sample size and four fish provided enough statistical power (*Power* = 1, *n* = 4, *f* = 1.71, *sig. level* = 0.05). A one-way ANOVA was used to assess the effect of temperature on S_crit_ and a Tukey’s HSD *post hoc* test was used to determine significant pair-wise comparisons between temperatures.

All statistical analyses were performed in R 3.4.3 [48]. Data were checked for assumptions of normality by the visual Q-Q norm plot and statistically with the Shapiro-Wilk test where *P* > 0.05 indicate normally distributed data. Assumptions of homogeneity were assessed using the Levene’s test where a *P* > 0.05 indicates homogeneity. Data that did not fit assumptions of normality were log-transformed prior to further statistical analysis. Data are presented as mean ± SE and results from statistical analyses are defined as significant at *P* < 0.05.

## Results

### Metabolic rates and aerobic scope

RMR increased significantly with temperature (Figs 2A and 2B) and there was a significant effect of weight and temperature*weight interaction on RMR (*P* < 0.05; Table 1). While the results for the two MMR methods differed considerably, temperature, weight and temperature*weight interaction all had a significant effect on MMR using either method (*P* < 0.05; Table 1). The chase MMR increased continuously with temperature, while flume MMR increased with temperature until ~27°C (Fig 2A & 2B). The MMR values from the flume were consistently higher across the temperature range than from the chase method, indicating that the metabolic rate reached during the chase likely was not the maximum possible for this species.

**Figure 2A and 2B.**
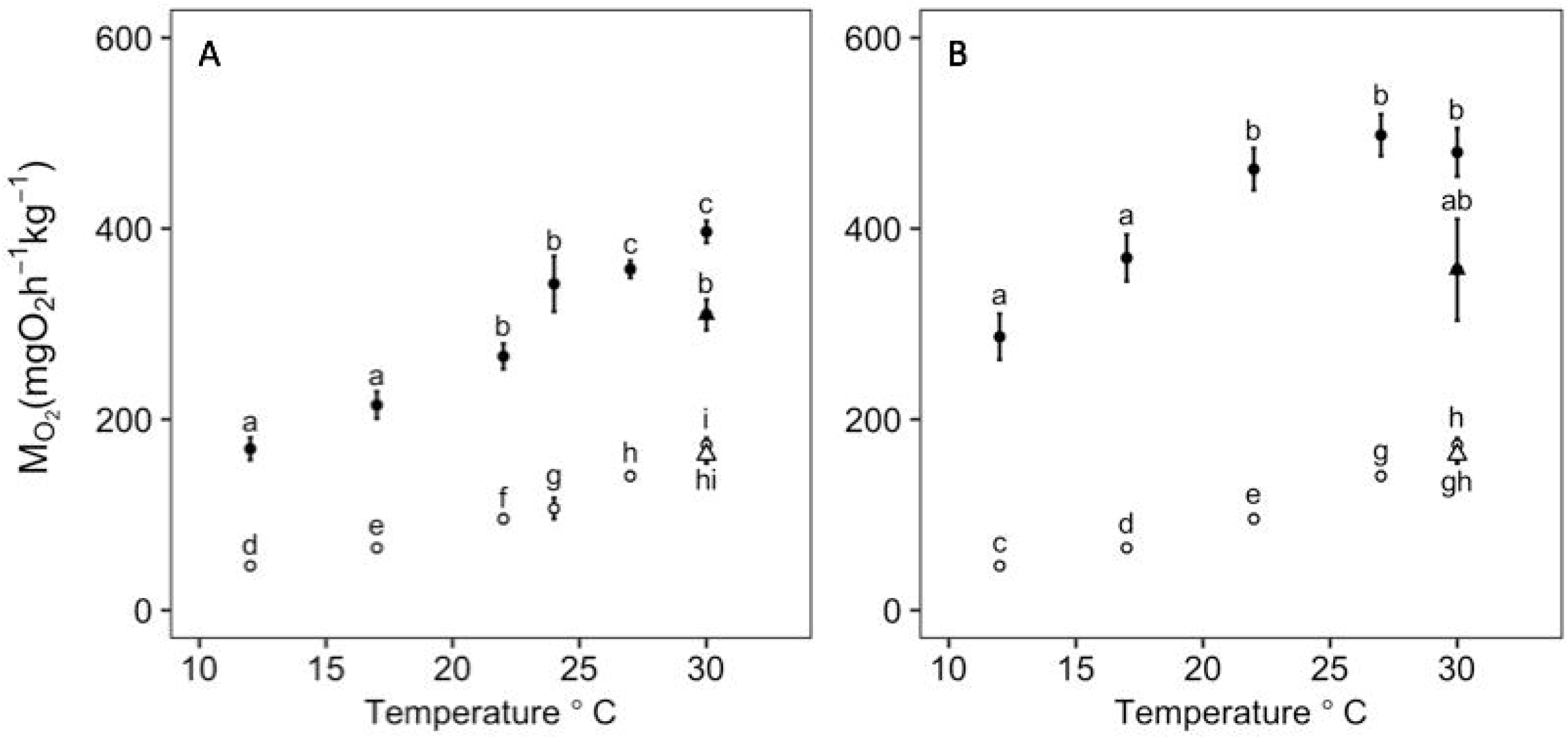
Effect of temperature on resting metabolic rate and maximum metabolic rate measured with a chase and a flume method. MMR (solid circles) and RMR (open circles) presented as mean ± s.e. normalized to a mean weight of 350g for each temperature treatment for chase method MMR (A) and flume method MMR (B). RMR is slightly different between (A) and (B) based on which fish were used for the respective MMR method. The 30_chronic_°C group is denoted by triangles. Tukey *post hoc* significance between treatments is shown by letters where data points with different letters indicate a significant difference (*P*<0.05).

**Table 1.** ANCOVA results for metabolic rates and aerobic scope.

| Variable | Effect | DF | F-value | P-value<sup>a</sup> |
| --- | --- | --- | --- | --- |
| AAS (chase) | Temperature | 6, 105 | 13.877 | < **0.001** |
|  | Weight | 1, 105 | 2.082 | > 0.05 |
|  | Temperature*weight | 6, 105 | 2.106 | 0.0586 |
| AAS(flume) | Temperature | 5, 48 | 6.185 | < **0.001** |
|  | Weight | 1, 48 | 6.599 | < **0.05** |
|  | Temperature*weight | 5, 48 | 4.033 | < **0.01** |
| MMR (chase) | Temperature | 6, 105 | 50.327 | < **0.001** |
|  | Weight | 1, 105 | 9.267 | < **0.01** |
|  | Temperature*weight | 6, 105 | 2.281 | < **0.05** |
| MMR (flume) | Temperature | 5, 48 | 16.244 | < **0.001** |
|  | Weight | 1, 48 | 8.927 | < **0.01** |
|  | Temperature*weight | 5, 48 | 3.147 | < **0.05** |
| RMR | Temperature | 6, 105 | 136.613 | < **0.001** |
|  | Weight | 1, 105 | 12.282 | < **0.001** |
|  | Temperature*weight | 6, 105 | 2.489 | < **0.05** |

AAS = absolute aerobic scope, MMR = maximum metabolic rate, RMR = resting metabolic rate
<sup>a</sup>P-values calculated from ANCOVA; bolded values are significant.

While the chase method did not achieve MMR, it still provided an estimate of submaximal exercise performance across a temperature range. The MMR using the chase method increased continuously with temperature and reached a maximum adjusted value of 396.65±11.48 mg O_2_ kg^−1^ hr^−1^at 30.0°C (the highest temperature measured; Table 2; Fig 2A). The MMR measured using the flume reached a maximum of 497.96±21.92 mgO_2_ kg^−1^ hr^−1^ at 27°C (Table 2; Fig 2B). The absolute aerobic scope using the flume method reached a maximum, typically referred to as “T_opt_” at ~24.4°C (Fig. 3B). There was a significant effect of temperature, weight, and the temperature*weight interaction on AAS (*P* < 0.05) using both MMR methods (Table 1). Using different MMR methods resulted in differences in the shape of the AAS curve (Fig 3A & 3B) and the estimated thermal optimum with consequences for its interpretation. All RMR, MMR and AAS values are reported in Table 2 and Q_10_ values are reported in Table 3.

**Table 2.** The MO_2adj_ mean ± S.E. values for all metabolic rates, aerobic scope and hypoxia tolerance.

| Temperature ($^\circ\text{C}$) | RMR | Chase MMR | Flume MMR | Chase AAS | Flume AAS | $S_{\text{crit}}$ |
| --- | --- | --- | --- | --- | --- | --- |
| 12 | $46.44 \pm 1.94$ | $169.12 \pm 11.78$ | $286.57 \pm 24.02$ | $117.67 \pm 7.55$ | $242.67 \pm 24.24$ | 19.65 |
| 17 | $65.27 \pm 3.27$ | $215.05 \pm 14.15$ | $369.27 \pm 24.47$ | $143.63 \pm 11.07$ | $303.55 \pm 24.70$ | 21.33 |
| 22 | $95.69 \pm 4.51$ | $266.03 \pm 13.32$ | $462.30 \pm 22.01$ | $167.18 \pm 12.13$ | $363.73 \pm 22.21$ | 21.80 |
| 24 | $106.61 \pm 11.03$ | $342.06 \pm 29.20^a$ | NA | $230.39 \pm 36.65^a$ | NA | NA |
| 27 | $140.55 \pm 4.48$ | $357.44 \pm 8.99$ | $497.96 \pm 21.92$ | $208.96 \pm 10.24$ | $351.65 \pm 22.12$ | 31.60 |
| 30 | $173.36 \pm 7.05$ | $396.65 \pm 11.48$ | $479.88 \pm 25.29$ | $213.20 \pm 13.33$ | $310.20 \pm 25.52$ | 37.88 |
| 30<sub>chronic</sub> | $163.14 \pm 9.28$ | $306.62 \pm 16.06$ | $356.82 \pm 53.05$ | $136.03 \pm 11.90$ | $198.33 \pm 53.54$ | 38.63 |

RMR = resting metabolic rate, MMR = maximum metabolic rate, AAS = absolute aerobic scope, $S_{\text{crit}}$ = critical %O<sub>2</sub> air saturation
<sup>a</sup>Adjusted $\text{MO}_2$ values for MMR and AAS at $24^\circ\text{C}$ are overestimated due to the average weight of fish in the $24^\circ\text{C}$ group to be smaller than the average weight for all study fish combined.

**Figure 3A and 3B.**
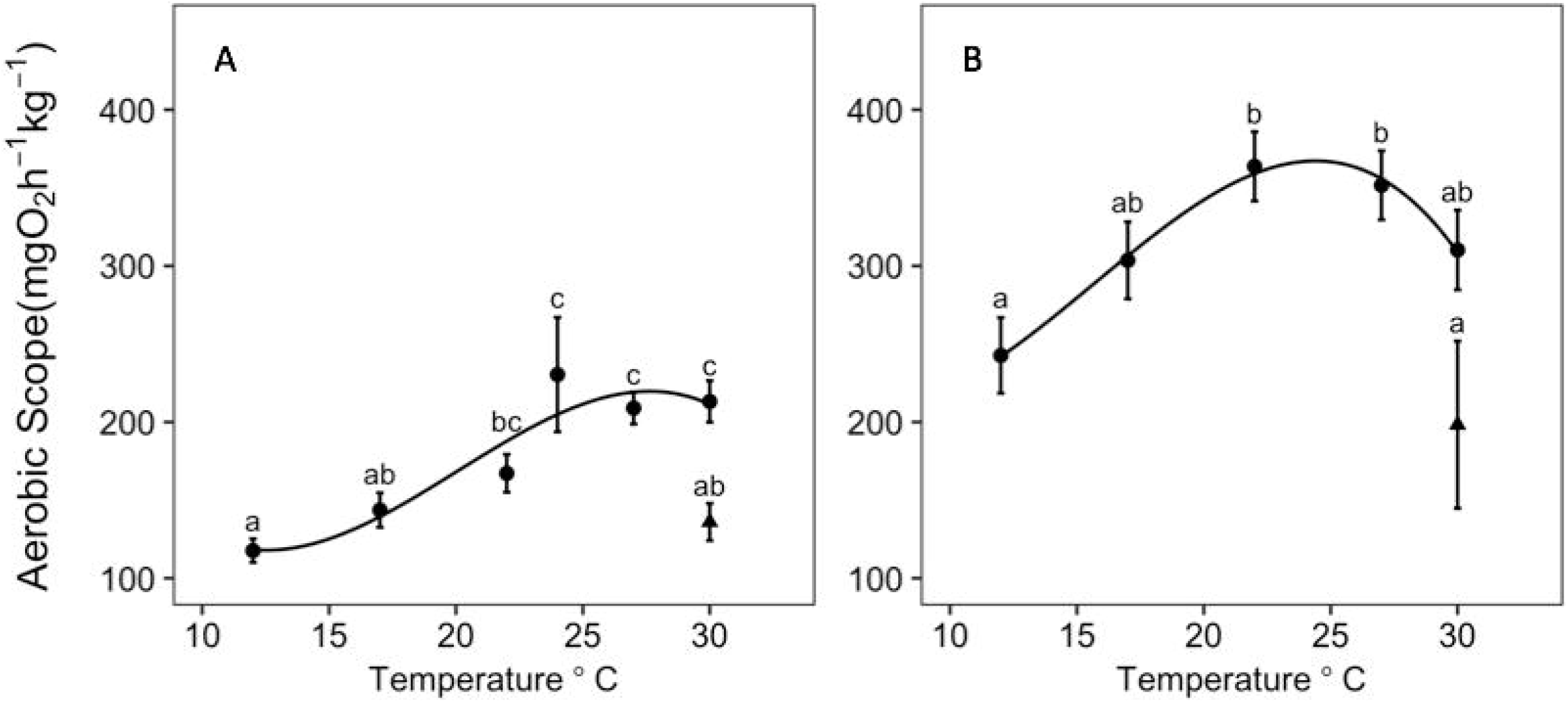
Effect of temperature on black sea bass aerobic scope. Aerobic scope (mean ± s.e.) of black sea bass normalized around a mean weight of 350g at each temperature treatment with the 30°C chronic group denoted by the black triangle. Letters indicate Tukey *post hoc* significance between groups where data points sharing a letter are not significantly different (*P*<0.05). Aerobic scope curves were generated from a) the chase MMR treatment (*y = 180.17 + 89.15x – 15.40x – 21.55x^3^; R^2^ = 0.878*) and b) flume MMR treatment (*y = 314.36 + 63.29x – 68.26x – 19.65x^3^; R^2^ = 0.994*).

**Table 3.** Q_10_ values separated between each temperature increment.

|  | 12-17°C | 17-22°C | 22-24°C | 22-27°C | 24-27°C | 27-30°C | 27-30°C<sub>c</sub> |
| --- | --- | --- | --- | --- | --- | --- | --- |
| AS<sub>chase</sub> | 1.49 | 1.35 | 4.97<sup>a</sup> | 1.56 | 0.72<sup>a</sup> | 1.07 | 0.24 |
| AS<sub>flume</sub> | 1.56 | 1.44 | NA | 0.93 | NA | 0.66 | 0.15 |
| MMR<sub>chase</sub> | 1.62 | 1.53 | 3.51<sup>a</sup> | 1.81 | 1.16<sup>a</sup> | 1.41 | 0.60 |
| MMR<sub>flume</sub> | 1.66 | 1.58 | NA | 1.16 | NA | 0.88 | 0.33 |
| RMR | 1.96 | 2.15 | 1.72 | 2.16 | 2.51 | 2.01 | 1.64 |

AAS = absolute aerobic scope, MMR = maximum metabolic rate, RMR = resting metabolic rate
<sup>a</sup>Slightly overestimated adjusted $MO_2$ for the 24°C fish is reflected in calculated $Q_{10}$ values.

### Critical %O_2_

The critical %O_2_ (S_crit_) increased significantly with increasing temperature (Fig 4; *F_5,18_* = 14.023, *P* < 0.05), directly correlated with RMR (Fig 5). There was no significant difference between 12°C (19.65 ± 1.72 %O_2_), 17°C (21.325 ± 1.75 %O_2_) and 22°C (21.80 ± 1.21 %O_2_), but S_crit_ increased significantly at 27°C (31.60 ± 1.67 %O_2_) and further at 30°C (37.875 ± 3.39 %O_2_). However, non-significance between 12, 17 and 22°C could be due to low sample size.

**Figure 4.**
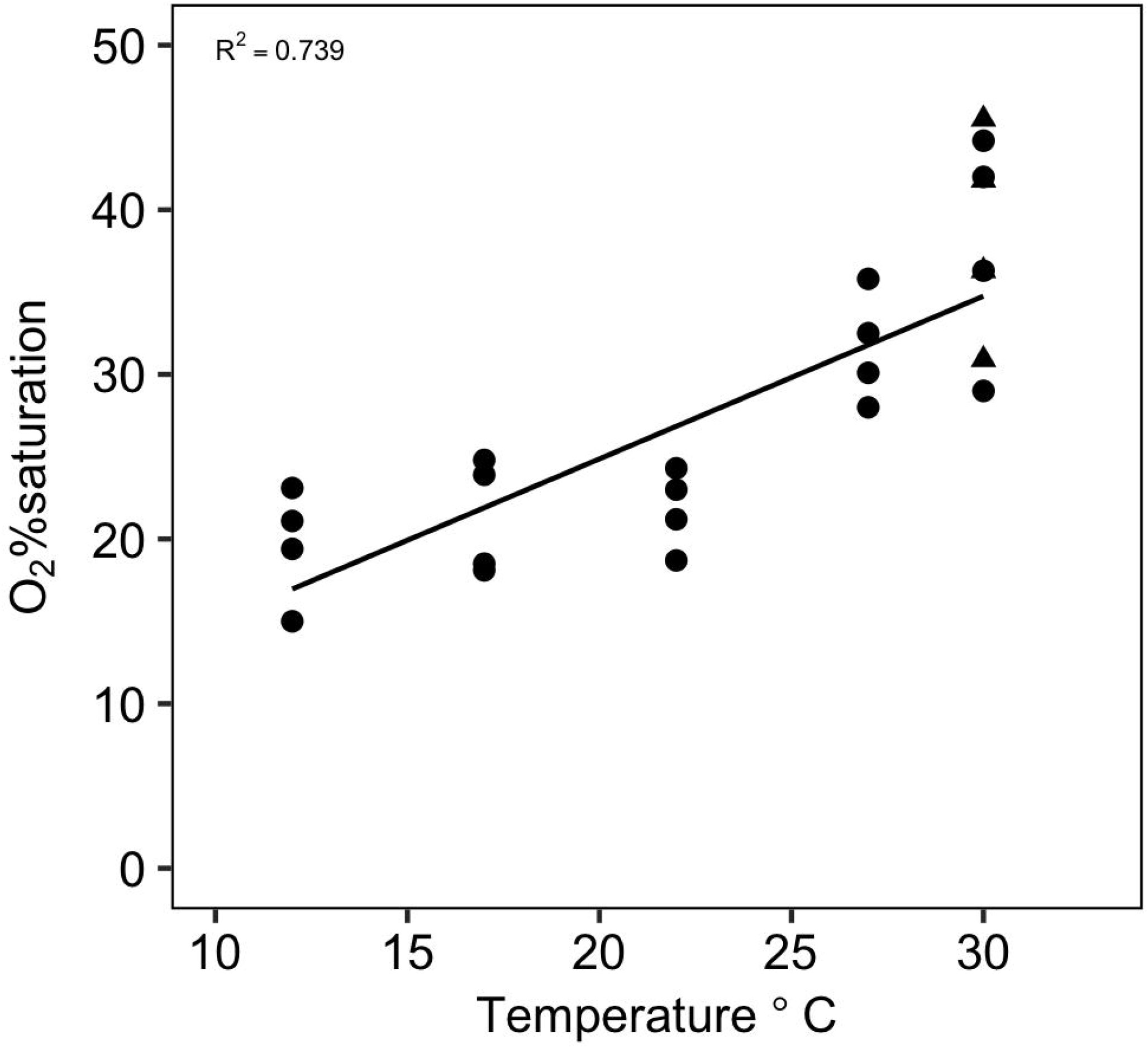
S_crit_ increases with increasing temperature. S_crit_ presented as %O_2_ air saturation for each temperature treatment. 30_chronie_°C treatment is denoted by a triangle and there is no significant difference between the 30_chronie_°C and acute 30°C treatments. A linear-regression was fitted for these data points (R_2_ = 0.793, *P*<0.001) showing an increase in S_crit_ (e.g. a decrease in hypoxia tolerance) with increasing temperature.

**Figure 5.**
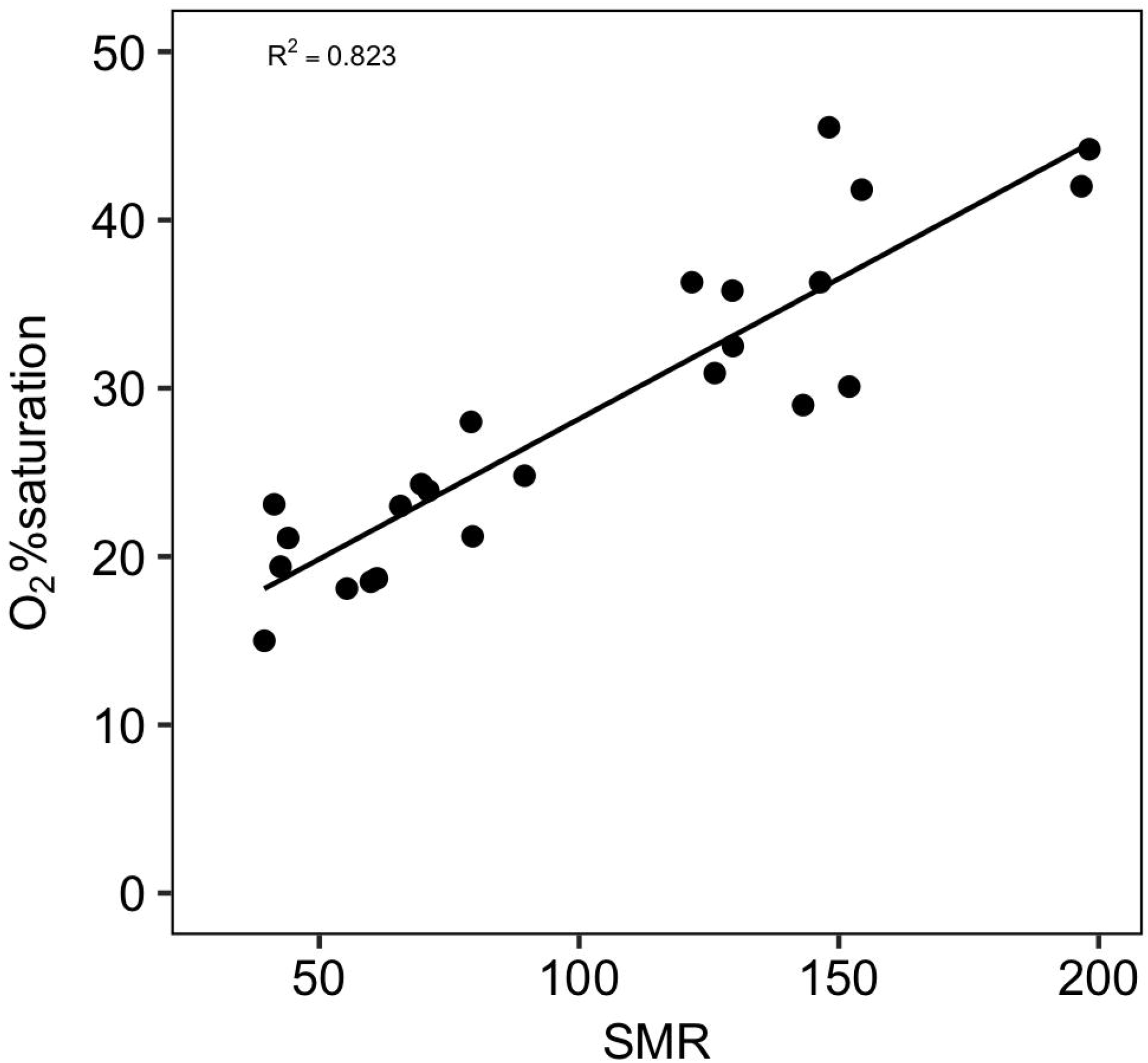
S_crit_ dependence on resting metabolic rate. S_crit_ is plotted against resting metabolic rate measured during the hypoxia experiment. A linear-regression was fitted for these data points (R_2_ = 0.823, *P*<0.001) and shows an increase in S_crit_ as metabolic rates also rise.

### Chronic high temperature exposure

The 30_chronic_°C group AAS using both MMR methods significantly decreased when compared to the 30°C treatment where fish were only held at this temperature for a week. Based on Tukey *post hoc* differences, RMR did not change significantly between the 30_chronic_°C and 30°C treatments but there was a significant decrease in MMR between the 30°C and 30_chronic_°C treatments. There was no significant difference in S_crit_ between 30°C and 30_chronic_°C treatments.

## Discussion

The primary objective of this study was to assess the use of physiological measurements to determine habitat suitability for the northern stock of black sea bass at current and future temperatures. We measured the oxygen consumption rate during two different exercise protocols. The flume yielded much higher metabolic rates, indicating that the chase method did not elicit MMR. Using the flume MMR, we found that AAS peaked at 24.4°C. S_crit_ increased with increasing temperatures as is typical of most (but not all, [49]) animals, including fishes [45]. Chronic exposure to 30°C resulted in a significant drop in AAS with no change in RMR or S_crit_. That S_crit_ increased with temperature in proportion to RMR, while MMR in the flume did not, suggests that chronic exposure to high temperature did not alter the capacity for oxygen uptake and transport, but that the capacity to generate ATP was reduced, perhaps due to a decrement in muscle function. The capacity for submaximal exercise (oxygen consumption following a chase to exhaustion) also increased across the entire temperature range further suggesting that the failure was not in the capacity for oxygen supply. Chronic exposure to 30°C led to further reductions in MMR using both methods, but no loss of oxygen supply capacity as estimated from S_crit_, suggesting continued deterioration in muscle function with longer exposure to warm temperatures.

Absolute AS typically increases with temperature up to a point, often termed “optimal”, and then declines at higher temperatures resulting in a roughly bell-shaped curve as has been identified in fishes that include, but is not limited to, juvenile European sea bass *Dicentrarchus labrax* [50], turbot *Scophthalmus maximus* [51], coho salmon *Oncorhynchus kisutch* [52], and sockeye salmon *Oncorhynchus nerka* [53]. However, some studies have found left- or right-skewed curves (e.g. [54]) while others find that AAS continues to increase up to the critical temperature for the species (i.e. no temperature optimum for AAS is identifiable; e.g. [55]). In our study, the black sea bass AAS curve was more bell-shaped with an estimated optimal temperature of 24.4°C. Bottom temperature in the southern portion of the black sea bass range typically hovers around 24-26°C during the summer ([56,57]; from U.S. East Coast Regional ESPreSSO model, [58]), which would suggest this area to be thermally optimal. However, if the loss in AAS at higher temperatures is due to a failure in muscular performance rather than potential oxygen supply, then 24°C may represent a maximum tolerable temperature rather than a temperature that allows optimal performance. In support of this interpretation, the Metabolic Index (which closely matches the factorial aerobic scope) declines with increasing temperature toward levels (~3 at 27°C in black sea bass) known to limit the geographic range of some species [17]. While the average bottom temperature in the southern portion of the northern stock of black sea bass is near 24°C during the summer months, there has still been a consistent expansion of their range northward into lower temperatures [59] further suggesting that the temperature eliciting maximum AAS is not, in fact, optimal. It is important to note that AAS is only a measured capacity to supply oxygen under maximum sustained exercise [21]. The required scope for other metabolic expenses (i.e. feeding, digestion; [60]) change with temperature in unknown ways and metabolic needs can change seasonally and with ontogeny [42]. Thus, AAS may in this case be an inappropriate predictor of fitness.

Black sea bass in the 30_chronic_°C treatment did not acclimate, indicated by no change in RMR or S_crit_ and a significant decrease in their MMR and AS. Norin et al. [55] similarly found that MMR and AAS in juvenile barramundi decreased significantly following 5 weeks at the highest study temperature (38°C). However, unlike black sea bass in our study, the juvenile barramundi RMR also decreased after the 5-week exposure. This same response has also been found for short-horn sculpin (*Myoxocephalus scorpius*) whose RMR was restored after being held at 16°C for 8 weeks to RMR values that were measured at 10°C [61]. The decrease in RMR can be an acclimation response to lower their energetic costs at high temperatures, but comes with its own caveats as this sometimes can reduce MMR. Importantly, black sea bass in the 30chronic°C treatment may have suffered stress from long-term captivity, which could also reduce AAS and time did not permit for a control chronic trial at a cooler temperature (although all fish were held for at least 5 days). Understanding the acclimation potential of black sea bass would benefit from future studies focusing on effects of a chronic treatment at each temperature tested.

S_crit_ decreased as temperature increased, most likely caused by rising RMR with higher temperatures, which has been shown in a majority of fish hypoxia studies (e.g. [23]; although see [49]). The 30_chronic_°C group did not have a significant decrease in hypoxia tolerance compared to the 30°C group, which agrees with no change in RMR between the two 30°C treatments. This suggests that the reduced MMR in 30_chronic_°C animals resulted from reduced capacity to generate ATP, rather than to supply oxygen. Black sea bass had lower S_crit_ than striped bass *Morone saxatilis* [62] and summer flounder *Paralichthys dentatus* [27], two important species found throughout the MAB that periodically experience hypoxic water during the summer months. However, when compared with fish that frequently experience hypoxia, such as largemouth bass and crucian carp [63] and juvenile barramundi [30], black sea bass were less hypoxia tolerant, especially in warmer water. Deutsch et al. [17] proposed a metabolic index (MI), as the ratio of oxygen supply to demand, which is effectively an estimate of a species’ time-averaged aerobic scope. By definition, the MI is equal to 1 at the S_crit_. A minimum MI of 2-5, indicating the capacity to supply oxygen at 2-5x the rate required at rest, is supportive of a population and delineates the equatorward distribution limit in the few species studied to date. Black sea bass factorial AS and MI both decreased with increasing temperature (Fig. 6). During the summer months when bottom water temperature is warmest along the coastal MAB, periodic hypoxic events occur after large phytoplankton blooms in the surface waters. In the past, these hypoxic events decreased bottom water PO_2_ below ~5.5kPa (26% saturation; 2.2 mg L^−1^ at 14°C; [35]), providing a metabolic index of ~1.3 at those temperatures for black sea bass. Such environments can be tolerated for short periods but are not likely supportive of a thriving population. At 30°C, even air-saturated water provides a MI of only 2.6 which is near the physiological limits of many species [17]. Therefore, when determining the suitable habitat, both temperature and oxygen must be taken into consideration as the interacting effects of these variables will effectively decrease optimal thermal habitat.

**Figure 6.**
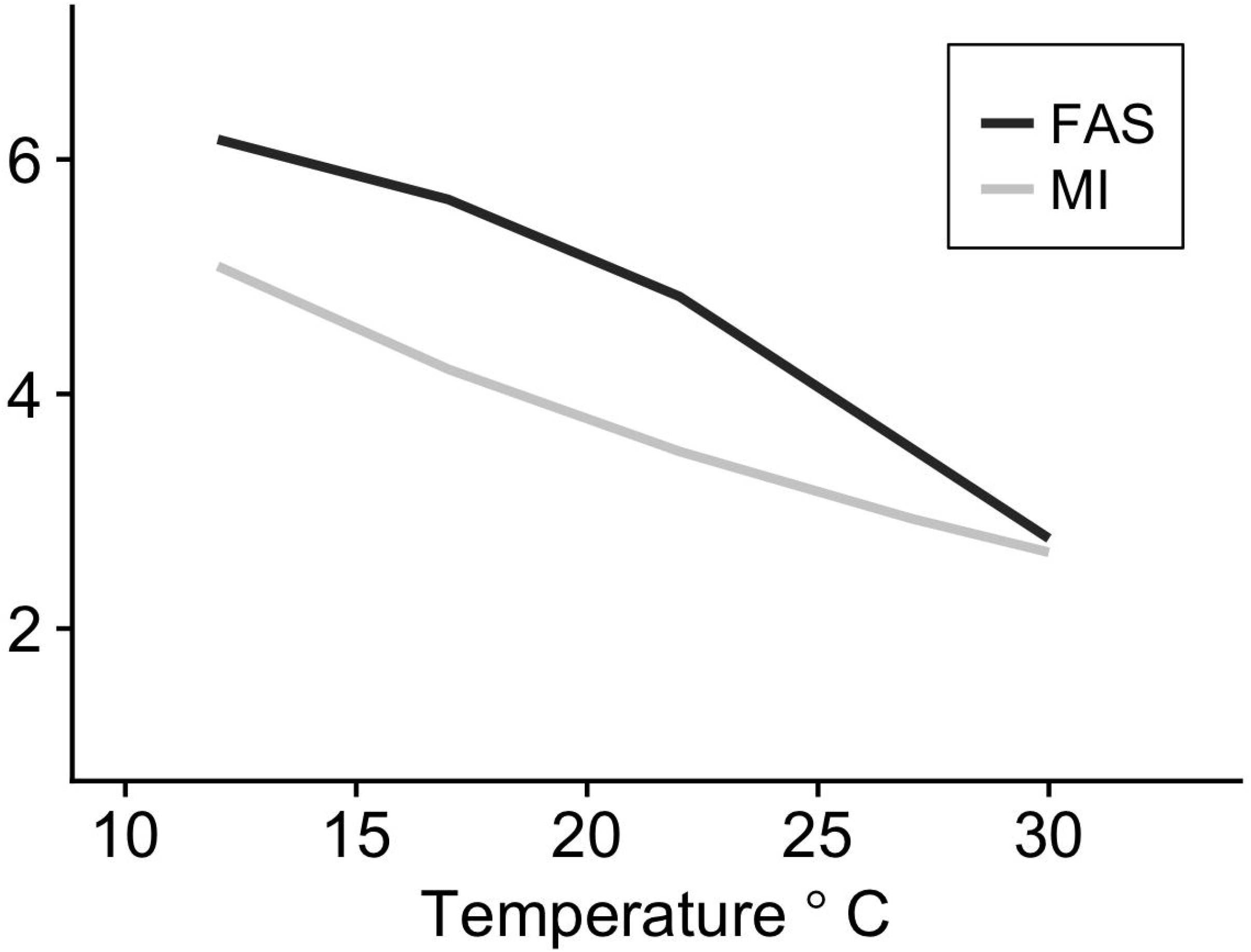
Factorial aerobic scope and metabolic index response to temperature. Factorial aerobic scope (FAS) and metabolic index (MI) plotted against temperature. Trends illustrate a decreasing trend in both measures as temperature increases. Both FAS and MI are unitless measures, but both measures scale similarly.

The chase method did not elicit MMR in black sea bass since MMR from the flume method was consistently higher. Which method, chase or flume, provides a more reliable measure of MMR and AAS is actively debated [64,65]. Whether a maximum rate of oxygen uptake is achieved by either method could depend on the type of swimming the study fish species naturally exhibits in the wild. Norin et al. [55] purposefully used a chase method for juvenile barramundi (*Lates calcarifer*), an ambush predator, that typically swims in quick bursts. In other cases, a fish will exhibit marked post-exercise oxygen consumption (EPOC; [66]), sometimes eliciting MMR minutes to hours after the cessation of exercise [67]. The swim flume method may be more appropriate for endurance swimming exhibited by pelagic fish such as tunas [65]. Different MMR methods may promote a certain type of swimming which could exhaust a fish before reaching MMR by depleting anaerobic stores, a noteworthy contributor to AAS [68]. For this study, we employed a sprint protocol for the swim-flume, which prompted similar burst swimming as in the chase method. However, during the chase protocol, black sea bass switched almost immediately to burst swimming accompanied with quick turning/flipping movements, compared to a slower transition and continuously straight burst swimming in the swim flume. The differences in MMR between the two methods could have been related to different swimming types, durations and/or speeds which could recruit more anaerobic resources [69] in the chase method, leading to exhaustion before reaching MMR.

In summary, the results from this study indicate that the northern stock of black sea bass reach a peak in AAS at ~24°C, which is warmer than in the northern portion of their range in the U.S. NES. The MI of 3.8 in air-saturated water, calculated from S_crit_ at 24°C, suggests relatively limited scope for sustained activity at that temperature [17]. We suggest that, rather than an optimal temperature, the peak in MMR and AAS indicates the maximum tolerable temperature, beyond which black sea bass experience a failure in some subcellular or organ systems that contribute to muscle performance. Our study only used individuals from the northern stock that were collected during the summer off of the New Jersey coastline. Metabolic research on the southern stock (south of Cape Hatteras, NC) and/or individuals from the northern stock in waters outside of New Jersey could reveal variation in some of these physiological metrics. However, the distribution of the northern stock of black sea bass has shifted northward [7] and this newly expanded habitat is almost 10°C colder than their apparent thermal optimum for AAS. We believe the preference for cooler waters reflects physiological limitation at higher temperatures, including possible limitation of oxygen supply relative to demand for growth and reproduction (reduced Metabolic Index) despite maintenance of oxygen supply capacity. However, many other factors, including food availability, additional energetic costs (e.g., evading predators, mating), or lower optimal temperatures for other critical processes may be important. This suggests AAS may not be the most appropriate predictor for habitat suitability in this species. Additionally, the northern stock of black sea bass population size has been increasing in the last decade [59], and this increase in biomass could be pushing part of the population northward. Regardless, the chronic exposure experiments presented here suggest little capacity for physiological adjustment to future temperatures. Black sea bass thermal habitat may shrink considerably in the southern region of the MAB as bottom water temperatures reach >27°C and continue to expand into the northern region of the MAB as ocean waters continue to warm, significantly impacting fisheries in these two regions.

## Acknowledgements

We thank Doug Zemeckis and Captain Chad Hacker (R/V Tagged Fish) for helping us collect black sea bass; Richard Brill and Andrij Horodysky for providing insight and suggestions for the study design; and the students at the Marine and Science Technology Academy for their assistance in animal husbandry. We also acknowledge the NOAA James J. Howard Laboratory personnel for their support and help throughout the study.

## Author Contributions

Conceptualization: VS, BS, GS

Data Curation: ES, BS, GS

Formal Analysis: ES

Funding Acquisition: VS, BS, GS

Investigation: ES, AA, RY, BP, JR, DW

Methodology: ES, AA, BS, JR, DW, GS

Project administration: ES, AA, VS, BS, GS

Resources: BS, BP, JR, DW, GS

Validation: ES, AA

Visualization: ES

Writing - Original Draft Preparation: ES

Writing - Review & Editing: ES, AA, RY, BS, VS, BP, JR, DW, GS

*The authors have declared that no competing interests exist.

## Supporting Information

**S1 Full Dataset. Experimental Parameters and Raw Data.**

**S1 File. Metadata Description.**

